# Comprehensive Sequencing of Environmental RNA from Japanese Medaka at Various Size Fractions and Comparison with Skin RNA

**DOI:** 10.1101/2024.09.23.614187

**Authors:** Kyoshiro Hiki, Toshiaki S. Jo

**Affiliations:** Health and Environmental Risk Division, National Institute for Environmental Studies, 16-2 Onogawa, Tsukuba, Ibaraki 305-8506, Japan; Research Fellow of Japan Society for the Promotion of Science, 5-3-1 Kojimachi, Chiyoda-ku, Tokyo 102-0083, Japan; Faculty of Advanced Science and Technology, Ryukoku University, 1-5 Yokotani, Oe-cho, Seta, Otsu, Shiga 520-2194, Japan

**Keywords:** environmental RNA, particle size distribution, biological monitoring, RNA-Seq, *Oryzias latipes*

## Abstract

Environmental RNA (eRNA) is emerging as a non-invasive tool for assessing the health of macro-organisms, but key information on its origin and particle sizes remains unclear. In this study, we performed comprehensive RNA-sequencing of eRNA (> 13 Gb/sample) collected from tank water containing Japanese medaka (*Oryzias latipes*), using sequential filtration through filters with pore sizes of 10, 3, and 0.4 μm. Fish skin RNA was also sequenced to reveal the origin of eRNA. Our results showed that the 3−10 μm fraction contained the lowest relative abundance of microbial RNA, the highest amount of medaka eRNA, and the largest number of detected medaka genes (5398 genes), while the 0.4−3 μm fraction had the fewest (972 genes). Only a small number of genes (42 genes) were unique to the 0.4−3 μm fraction. These findings suggest that a 3 μm filter is optimal for eRNA analysis, as it allows for larger filtration volumes while maintaining the relative abundance of macro-organism eRNA. Furthermore, 81% of the genes detected in eRNA overlapped with skin RNA, indicating skin is a major source of fish eRNA.

**Synopsis:** The 3 µm filter is recommended to maximize the detection of eRNA from macro-organisms while reducing the relative abundance of microbial RNA.

**TOC Art:** 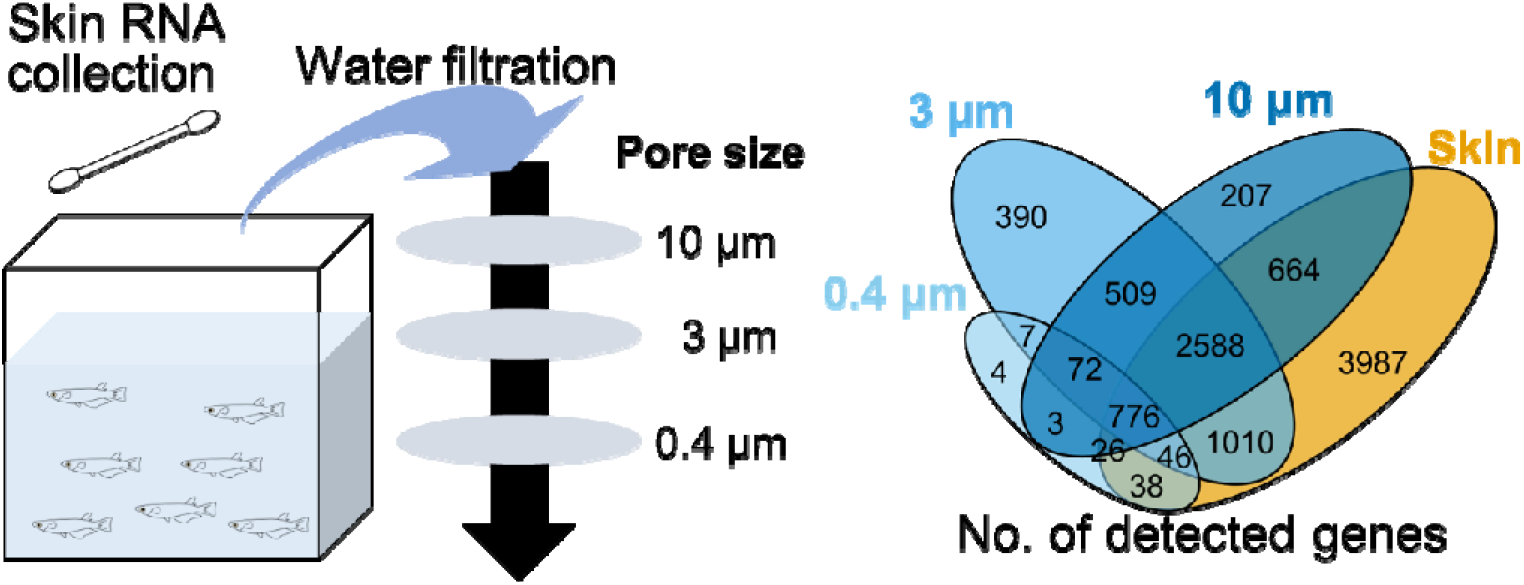

## 1. INTRODUCTION

Environmental RNA (eRNA) analysis, emerging as a complementary tool to environmental DNA (eDNA) analysis, has significant potential for advancing our ecological monitoring (Cristescu 2019; Tsuri et al. 2021; Veilleux et al. 2021). Both of the molecular tools detect DNA and RNA fragments released from macro-organisms such as fish in environmental samples (i.e., eDNA and eRNA) to infer species’ occurrence and composition rapidly and non-invasively without traditional capture-based surveys (Veilleux et al. 2021; Yao et al. 2022). While DNA is responsible for storing and transferring genetic information, RNA transmits the genetic code for protein creation and is synthesized from DNA when required. Owing to the RNA characteristics, eRNA analysis can provide the physiological information (e.g., reproduction, development, stress response, and living/dead) that cannot be inferred by eDNA analysis (Yates et al. 2021; Stevens and Parsley 2023; Parsley and Goldberg 2024). In particular, eRNA expression profiling by high-throughput sequencing has allowed for the detection of a variety of genes that are up-/down-regulated in response to environmental stressors like chemical exposure and increased temperature (Hechler et al. 2023; Hiki et al. 2023), suggesting the potential of eRNA analysis as a non-invasive tool for assessing the real-time physiological status of organisms.

Despite its potential, the current eRNA applications for assessing the physiological status are at the stage of the proof-of-concept (e.g., aquarium experiment) and have several limitations against its practical applications to field environments (Yates et al. 2021; Hiki et al. 2023). First, RNA extracted from an environmental sample contains not only extra-organismal RNA released from macro-organisms but also organismal RNA of microbes. According to the previous studies, the most of sequenced eRNA reads (> 99%) from aquarium samples were derived from bacteria and fungi and only a small proportion of the macrobial eRNA reads derived from fish or zooplankton were available (Hechler et al. 2023; Hiki et al. 2023). This point brings to the substantial inefficiency in expression profiling of macrobial eRNA especially using high-throughput sequencing without any pretreatments such as PCR and removal of microbial RNA. Even when the PCR-based detection approaches (species-specific or metabarcoding) are used, detectability of eRNA derived from nuclear mRNA genes is much lower than those derived from rRNA and mitochondrial genes (Marshall et al. 2021; Jo 2023), which would be further problematic particularly in a turbid water environment where suspended particle matter and PCR inhibitors (e.g., humic and tannic acids, proteins, and metal ions) are abundant (Robson et al. 2016; Kumar et al. 2022).

Second, not all genes expressed within an organism may be detected as eRNA. Macro-organismal eDNA and eRNA (collectively, environmental nucleic acids [eNAs]) is considered to be derived mainly from epidermis, mucus, gills, feces, and gametes (Merkes et al. 2014; Barnes and Turner 2016; Tsuri et al. 2021; Wu et al. 2024). On the other hand, unless an organism is dead or damaged, internal organs including brain and heart, nerve cells, bones, and muscles are unlikely to be released so much from the individual and be the major origin of eNAs. Previous eRNA studies reported that the numbers of genes detected in whole body of organisms were overwhelmingly abundant compared with those detected from the surrounding water (Hechler et al. 2023; Hiki et al. 2023). The results could be attributed to the biased origin of eRNA particles, as well as its shorter persistence (Marshall et al. 2021; Jo et al. 2023), suggesting that some types of gene within an organism can be never inferred by eRNA analysis. In addition, even if a target gene is detected from both organisms and surrounding environments, its expression patterns might be different between in organisms and in the environments (Hiki et al. 2024). Nevertheless, for a proper validation of its monitoring performance, it is necessary to understand what kinds of genes are released into the environment and what kinds of tissues and organs the genes are derived from.

To address the knowledge gaps, we conducted an aquarium experiment using Japanese medaka *Oryzias latipes*, a model species of vertebrates. Rearing water was sequentially filtered through different pore size filters (10, 3, and 0.4 µm) to obtain size-fractioned eRNA samples (i.e., 0.4−3, 3−10, and >10 µm size fractions) that were comprehensively sequenced. In our previous experiment using a quantitative real-time PCR (qPCR), zebrafish *Danio rerio* eRNA of mitochondrial cytochrome b was abundantly present in 3−10 µm size fraction (57.6 % of all fractions on average; Jo, 2024). We hypothesized that 3−10 µm fraction would yield the largest number of detected genes. If confirmed, this hypothesis could enhance the efficiency of fish eRNA collection by optimizing filter pore size and increasing the proportion of fish eRNA compared with microbial RNA. In addition, we also collected the skin RNA, one of the possible major eNA origin, from *O. latipes* individuals reared in our experiment. The skin RNA samples were also comprehensively sequenced to compare transcriptome profiles between water and skin and to understand the contribution of skin-derived genes to eRNA profiles.

## 2. MATERIALS AND METHODS

### 2.1. Test Organisms

Japanese medaka *O. latipes* was obtained from a brood stock maintained for more than 15 years at National Institute for Environmental Studies, Japan (NIES-R strain), as described in our previous study (Hiki et al. 2023). Briefly, the brood stock was kept in 5 L glass tanks with dechlorinated tap water from Tsukuba (hardness: about 80 CaCO_3_ mg/L) at 25 °C under under light-dark cycles of 16:8 hours. Fertilized eggs were collected, and the hatched larvae were reared under identical conditions, feeding on freshly desalinated brine shrimp (*Artemia* spp.) nauplii until use in experiments.

### 2.2. eRNA and skin RNA collection

Thirty-five 27-days-old medaka fish were transferred to a glass tank (8 cm × 19 cm × 20 cm height) containing 2 L of dechlorinated tap water, with four replicates (i.e., a total of 140 organisms). Twelve additional fish from the same brood stock were used for measuring body length and wet weight (21.2 ± 2.3 mm and 13.2 ± 0.5 mg, mean ± SD). The tanks were kept at 25 °C under light-dark cycles of 16:8 hours for 96 hours without aeration and feeding. The measured water temperature, pH, conductivity, and dissolved oxygen concentration were 24.5 °C, 7.7, 34.8 mS/m, and 7.65 mg/L at the end.

After 96 hours, water samples ranging from 750 mL to 1250 mL were sequentially filtered (Table S1) using polycarbonate membranes with nominal pore sizes of 10 (large), 3 (middle), and 0.4 μm (fine) (Millipore, USA) and filter funnels (Magnetic Filter Funnel, 500 ml capacity, Pall Corporation, USA) connected with silicon tubes (SR-1554, internal diameter: 5 mm) and aspirated by a pump. As a blank control, dechlorinated tap water was similarly filtered. The polycarbonate membrane was cut in half with scissors and then both pieces were immediately used for RNA extraction.

To investigate eRNA sources, fish skin RNA was analyzed. Due to the impracticality of collecting feces from the small medaka used in this study, feces as another potential eRNA source were not analyzed. Following the method of a previous study (Breacker et al. 2017), skin RNA from eight medaka fish after 96 hours was collected using a cotton swab while the fish were held in an aquarium net on an aluminum foil. This procedure was repeated for a total of 16 randomly selected fish from each tank.

### 2.3. RNA extraction

eRNA extraction was performed using RNeasy Mini Plus Kit (Qiagen, Germany). Six hundred μL of RLT buffer containing β-mercaptoehanol was added to the polycarbonate membrane or swab tip placed in a 1.5 mL tube. The tube was then gently rotated using a rotator (RT-50, Taitec, Japan) for 1.5 hours at room temperature. Then the buffer was combined from two tubes, corresponding to one polycarbonate membrane and two swab tips (i.e., representing 16 fish). The RNA extraction was performed following the manufacturer’s protocol, but with an additional step of DNase digestion on the RNeasy column membrane using RNase-free DNase set (Qiagen). This digestion was performed for 30 minutes following the wash with RW1 buffer and was repeated twice to ensure complete DNA removal. The RNA was then eluted in 50 µL of water. The RNA integrity number equivalent (RINe) and concentration were determined using 4200 TapeStation (Agilent Technologies, USA).

### 2.4. RNA sequencing

A cDNA library was constructed using SMART-Seq v4 Ultra Low Input RNA Kit (Illumina,USA) following the manufacturer’s protocol. Note that this library construction method relies on 3′ oligo(dT)-primed cDNA synthesis, which may introduce a bias toward 3′ regions, particularly in degraded eRNA samples. The constructed library was sequenced on NovaSeq X Plus using paired-end 101 bp reads by Macrogen Japan (Tokyo), achieving 135−185 million reads for each sample. Q30, a ratio of bases that had phred quality score of over 30, exceeded 92% for all samples. All raw sequencing reads were deposited on the Sequence Read Archive (accession number: DRR594069 −DRR594084 under the project PRJDB15208).

### 2.5. Data analysis

Data analysis was performed following the procedure described in our previous study (Hiki et al. 2023). The sequenced reads were mapped to the reference genome of *O. latipes* (accession number: GCF_002234675.1) using HISAT2 ver. 2.2.1 (Kim et al. 2019) with default settings. Quality trimming was not performed as the sequenced reads for all samples were of high enough quality. The mapped reads were counted using the GTF file and HTSeq ver. 0.11.3 (Anders et al. 2015) with the “-s reverse” option, and then analyzed using R software ver. 4.3.1 and edgeR ver. 4.2.1 (Chen et al. 2024). Genes with low read counts were removed to keep genes that have count-per-million (CPM) above 10 in 70% of each treatment samples (i.e, in at least 3 samples). Gene set enrichment analysis (GSEA) was performed using ShinyGO ver. 0.80 (Ge et al. 2020) with all available databases (e.g., Kyoto Encyclopedia of Genes and Genomes, Gene Ontology). The cutoff for significance in GSEA was set at a false discovery rate (FDR)-adjusted p-value of 0.05. Transcript integrity number (TIN) (Wang et al. 2016), as an index for mRNA integrity at the transcript level based on mapped read coverage, was assessed using RSeQC ver. 2.6.4 (Wang et al. 2012). To evaluate whether the sequencing depth was sufficient for gene detection and expression analysis in eRNA, a Monte Carlo simulation was performed using the “sample_n” function in the dplyr R package. In this simulation, reads mapped to the reference genome were randomly selected, and the number of genes identified by the selected reads was counted. The bivariate relationships of mean logarithmic CPM between sample types (i.e., 0.4 μm filter, 3 μm filter, 10 μm filter, and skin) were analyzed by standardized major axis (SMA) regression using the smatr R package ver. 3.4-8 (Warton et al. 2012). Principal component analysis (PCA) was conducted based on logarithmic CPM of eRNA samples using the prcomp R function with the “scale = TRUE” option. An example R code can be found on GitHub (https://github.com/KyoHiki/eRNA_filter_analysis).

To determine biological sources of unmapped reads, we performed taxonomic identification using DecontaMiner ver. 1.4 (Sangiovanni et al. 2019) with the database provided by the software’s web site (https://github.com/amarinderthind/decontaminer).

### 2.6. Quality assurance and quality control

Prior to the laboratory experiments, all equipment including glass tanks and filter funnels were treated with 10% bleach solution and thoroughly rinsed with Milli-Q water to remove DNA and RNA (Jo et al. 2023). All experiments were performed while wearing disposable gloves to minimize contamination. All equipment for molecular analysis (e.g., pipettes and Laminar flow hood) was decontaminated by ultraviolet irradiation before use. The DNA contamination was checked in both blank control and eRNA samples by conventional PCR using primers PG17 for DMY gene (Matsuda et al. 2002) and 1% agarose gel electrophoresis.

## 3. RESULTS AND DISCUSSION

### 3.1. Overview of extracted RNA and RNA sequencing

The RINe ranged from 4.7 to 8.0 for all RNA samples, with an average of 6.4 ± 1.1 (*n* = 16) (Table 1). The RINe was significantly lower in eRNA collected with 3µm filters (mean: 5.1 ± 0.5) than 0.4 µm filters (mean: 7.2 ± 0.8) and skin RNA (mean: 6.8 ± 0.6) (two-sided Tukey’s test, *p* < 0.05). RNA concentrations ranged from 2.0 to 19 ng/µL for eRNA and 3.2 to 13 ng/µL for skin RNA. The concentration of eRNA collected with 0.4 µm filters (16.4 ± 2.4 ng/µL) was significantly higher than that with larger pore sizes (two-sided Tukey’s test, *p* < 0.05).

**Table 1.** Summary of extracted RNA and RNA sequencing.

|  | eRNA captured with filters |  |  | Skin RNA |
| --- | --- | --- | --- | --- |
| | 10 $\mu\text{m}$ | 3 $\mu\text{m}$ | 0.4 $\mu\text{m}$ | |
| RNA integrity number equivalent (RINe) | $6.4 \pm 1.0$ | $5.1 \pm 0.5$ | $7.2 \pm 0.8$ | $6.8 \pm 0.6$ |
| RNA concentration (ng/$\mu\text{L}$) | $4.2 \pm 1.5$ | $3.7 \pm 1.5$ | $16.4 \pm 2.4$ | $8.2 \pm 4.0$ |
| Number of sequenced reads (million) | $174 \pm 12$ | $173 \pm 14$ | $164 \pm 13$ | $152 \pm 11$ |
| Ratio of mapped reads (%) | $0.6 \pm 0.1$ | $1.6 \pm 1.1$ | $0.3 \pm 0.0$ | $80.9 \pm 5.8$ |
| Transcript integrity number (TIN) for mitochondrial genes | 85 (83–88) | 87 (87–90) | 86 (84–87) | 92 (90–94) |
| TIN for nuclear genes | 14 (6–27) | 23 (13–37) | 10 (0–24) | 82 (75–86) |
| Number of detected genes | 4845 | 5398 | 972 | 9135 |

Mean ± standard deviation ( $n = 4$ ) except median and interquartile range (25 and 75 percentiles) for TIN of each gene.

For sixteen libraries, a total of about 2.65 billion reads were obtained with a length of 101 base pairs (see Tables 1 and S2). Of the sequenced reads, about 81% of skin RNA were mapped to the reference genome, while less than 3.0% of RNA captured with filters were mapped. This low mapping ratio for eRNA is consistent with previous observations for medaka (Hiki et al. 2023) and *D. pulex* (Hechler et al. 2023), indicating that the majority of

RNA reads were derived from nontarget microorganisms, such as bacteria and fungi. The detailed composition of microorganisms is shown in Fig S1. Although not statistically significant (*p* > 0.05, Tukey’s test), the mapping ratio was highest for eRNA captured with 3 μm filters (1.6 ± 1.1%), followed by 10 μm filters (0.6 ± 0.1%), and lowest for 0.4 μm filters (0.3 ± 0.0%). The higher ratios for 3 μm and 10 μm filters are reasonable, as non-target microorganisms can pass through these filters while cells or organelles may be retained (Turner et al. 2014; Jo 2024).

TIN values were sufficiently high in skin RNA for both mitochondrial and nuclear genes with medians ranging from 75 to 94 (Figure 1, Table 1). In contrast, TIN values of eRNA were generally low for nuclear genes, with the medians < 25, but high for mitochondrial genes (median TIN: 83−90). These results are consistent with our previous findings (Hiki et al. 2023). Additionally, TIN values varied slightly among eRNA samples filtered through different pore sizes: 3 μm ≥ 10 μm > 0.4 μm for nuclear genes. The reasons for lower TIN values observed with the 0.4 μm filter is unclear, but it might be partly due to RNA in this fraction not being protected by membranes, cell debris, or RNase-inhibitor (e.g., free fatty acids (Inoue et al. 2022)), making it more susceptible to degradation. Note that TIN values in eRNA were not consistent with RINe values. This discrepancy is because RINe reflects the overall integrity of the RNA collected, which predominantly consists of RNA from non-target microorganisms, whereas TIN specifically measures the integrity of medaka mRNA.

**Figure 1.**
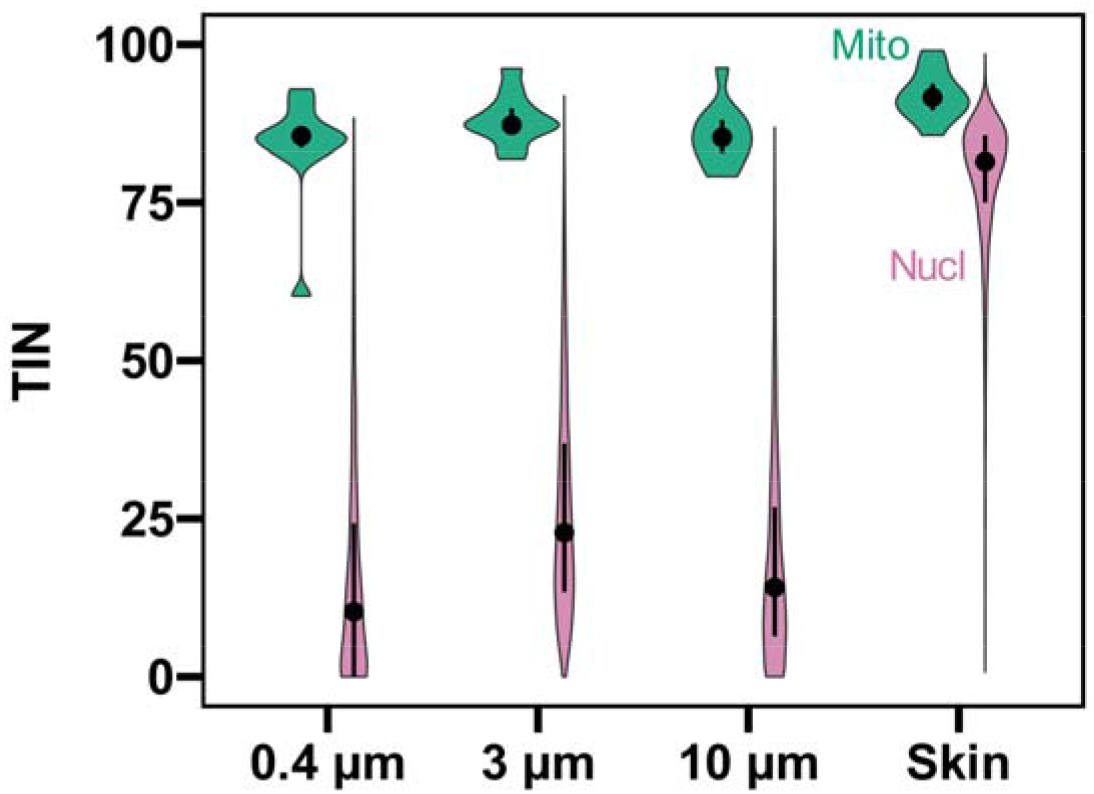
Distribution of transcript integrity number (TIN) in Japanese medaka eRNA captured with filters of three different pore sizes and in fish skin RNA. Different colors represent mitochondrial (green) and nuclear (pink) genes. The dot represents the median for detected genes, and the bar indicates the interquartile range (25th and 75th percentiles).

The number of detected genes was highest for skin RNA (9153), followed by 3 μm filter (5398), 10 μm filter (4845), and lowest for 0.4 μm filter (972). Although these values might not represent the maximum number of genes detectable in each fraction, Monte Carlo simulations demonstrated that the number of detected genes in all sample types was close to a plateau (Figure S1), indicating that the number obtained was close to the number of genes presented in each fraction. However, the number of genes with > 5 read counts did not reach a plateau in eRNA samples likely due to the sequencing reads being consumed by microbial RNA, whereas it did in skin RNA samples. This suggests that the sequencing depth for the medaka transcriptome in this study (approximately 2 million reads even for 3 μm filter, Table S2) was not sufficient for quantitative analysis of eRNA. Typically, >40−80 million reads are required to quantify lowly expressed genes and to perform differential expression analyses (Sims et al. 2014). Therefore, caution should be exercised in subsequent quantitative analyses for eRNA.

### 3.2. Filter eRNA versus skin RNA

About half of detected genes in fish skin RNA (56%) were also detected in water as eRNA (Figures 2A and 2B). Genes that were detected in both water and skin had higher expression levels (median log_2_ CPM: 5.4) compared to those detected only in skin RNA (median log_2_ CPM: 4.5) (Figure 2C), whereas TIN values were not different between them (median TIN: 82 for both). Logarithmic CPM in skin RNA correlated with that in eRNA (Figure 2E). GSEA showed the significant enrichment of pathways related to endomembrane system (GO: 0012505), Golgi apparatus (GO: 0005794), vesicle (GO:0031982), endosome (GO: 0005768), and bounding membrane of organelle (GO: 0098588) in gene sets detected both in water and skin (Figure 2G, FDR-adjusted p-value < 0.05). In contrast, genes detected only in fish skin RNA were enriched in pathways related to mitochondrion (GO: 0005739), ncRNA processing (GO: 0034470), tRNA processing (GO: 0008033), and extracellular region (GO: 0005576) (Figure 2F). These findings suggest that eRNA in water originates significantly from fish skin RNA, particularly from those that are highly expressed or actively secreted by endomembrane systems such as endosomes and the Golgi apparatus.

**Figure 2.**
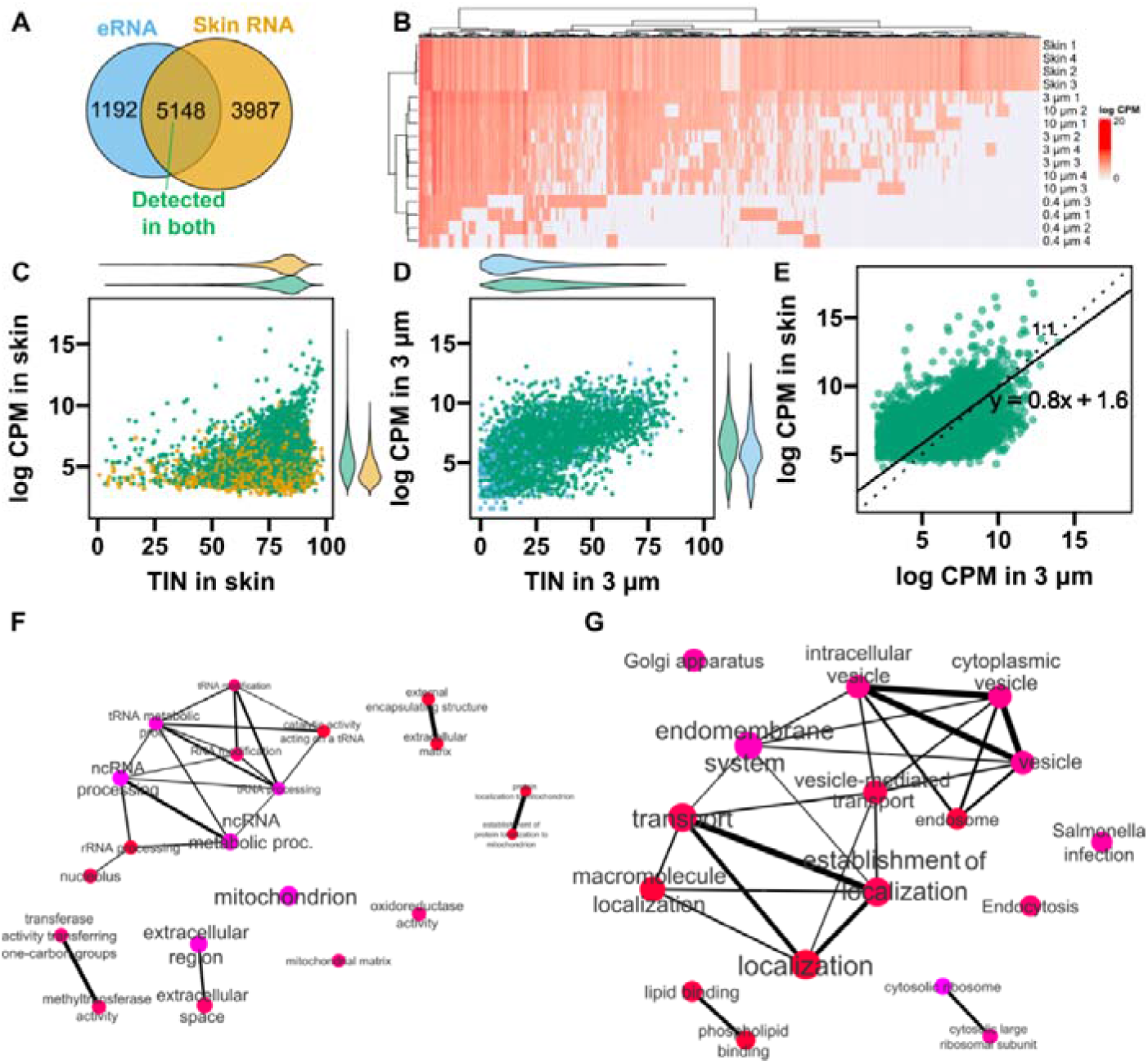
Comparison of expression profiles between fish skin and environmental RNA (eRNA). (A) Venn diagram showing the overlap of detected genes between fish skin RNA and eRNA in water. (B) Heatmap of all the detected genes. Color represents logarithmic (log_2_) counts per million reads (CPM). Number of a sample name represent the ID of water tank. (C) Relation between TIN and logarithmic CPM in fish skin RNA. Green indicates genes detected in both eRNA and skin RNA (“detected in both”), while yellow rsssepresents genes specifically detected in fish skin. Mean logarithmic CPM for each gene was calculated from four control samples. (D) Relation between TIN and logarithmic CPM in eRNA collected by the 3 µm filter. Light blue indicates the genes specifically detected in eRNA. (E) Comparison of logarithmic CPM between eRNA collected by the 3 µm filter and skin RNA. Solid and dotted lines represent standardized major axis regression and 1:1 line, respectively. (F) Pathway networks enriched in skin-specific RNA (3987 genes) compared to all genes detected in skin RNA. Vivid color and larger nodes are more significantly enriched (FDR-adjusted p-value < 0.05) and larger gene sets, respectively. Edge widths represent the degree of gene overlap. (G) Pathway networks enriched in genes “detected in both” (5148 genes) compared to all genes detected in skin RNA.

Skin and skin mucus serve as essential barriers that protect fish from pathogens. The endomembrane system in fish skin is crucial for immune responses, including antigen processing, antibody production, pathogen destruction, and immune signaling (Zhu et al. 2013; Brinchmann 2016; Smith et al. 2019). Therefore, our findings that endomembrane-related genes were released from skin into water suggests the potential of eRNA as a non-invasive assessment tool for immune function. Previous studies found immune-related genes from fish epidermal mucus (Greer et al. 2019; Andrzejczyk et al. 2022), further highlighting this potential. This potential is also supported by the detection of immune-related genes (e.g., interleukin [il17rc, il17rd, il6st, and il7r], nf-κβ [nfkb1 and nfkb2]) in both eRNA and skin RNA and by the enrichment of *Salmonella* infection pathway (Path:ola05132) in gene sets detected both in water and skin (Figure 2G).

Although most of detected genes in eRNA captured with filters (81%) were detected in fish skin (Figure 2A), the remaining 19% were not detected in skin, suggesting that eRNA originates from multiple sources, such as urine and feces, in addition to skin. GESA revealed that genes specifically detected in eRNA, including semaphorin and troponin gene families, were enriched in pathways related to extracellular region (GO: 0005576), regulation of axonogenesis (GO: 0050770), and myofilament (GO: 0036379), compared to all the genes detected in eRNA (6340 genes) (Figure S3). Genes specifically detected in eRNA had lower TIN values compared to those detected in both, except for eRNA collected using the 0.4 μm filter (Figures 2D and S4). This suggest that genes originating from sources other than skin (e.g., feces and urine) may be more degraded, resulting in lower CPM values.

### 3.3. eRNA at different size fractions

As described above, the numbers of genes for 3 µm and 10 µm filters were similar and higher than that for 0.4 μm filter. Additionally, most of genes detected by 0.4 µm filter were also detected by 3 µm and 10 µm filters (Figure 3A), with only 4.3% (42 genes) being unique to 0.4 µm filter. Although variation was large, logarithmic CPM correlated well across the fractions (*p* < 0.001), in particular between 3 µm and 10 µm (Pearson’s r = 0.89), and the 95% confidence intervals of SMA regression slope and intercept included 1.0 and 0.0, respectively (Figure 3C−E). PCA analysis also showed that 3 µm and 10 µm filters clustered closely together, separating distinctly from 0.4 µm filter (Figure 3F).

**Figure 3.**
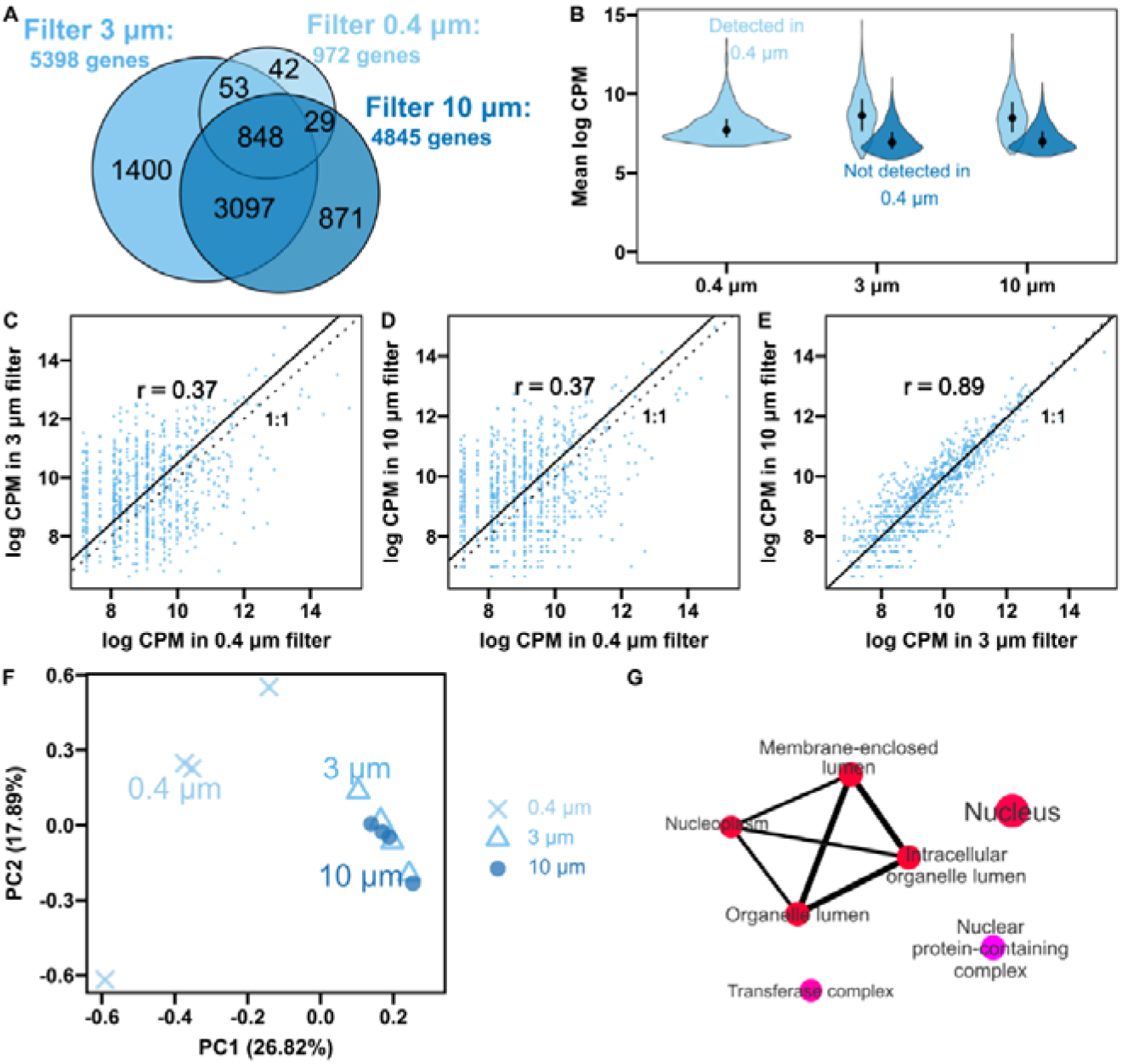
Profiles of Environmental RNA (eRNA) captured by filters with three different pore sizes. (A) Venn diagram showing the overlap of detected genes among eRNA captured with 3 µm, 10 µm, and 0.4 µm filters. (B) Violin plots showing the distribution of mean log CPM for eRNA across different filters. Error bars indicate the median and interquartile range (25 and 75 percentiles). Light blue represents genes detected in 0.4 µm filter, while blue represents genes not detected in 0.4 µm filter. (C−E) Scatter plots comparing log CPM values between pairs of filters, with Pearson’s r values: 0.4 µm vs. 3 µm (C), 0.4 µm vs. 10 µm (D), and 3 µm vs. 10 µm (E). Dotted and solid lines indicate 1:1 ratio and standardized major axis regression, respectively. (F) Principal component analysis based on logarithmic CPM of commonly detected genes. PC1 and PC2 explained 26.82 and 17.89% of the data variance. Different symbols represent different filter sizes. (G) Pathway networks of cellular component enriched in genes that were not detected in 0.4 µm filter (5368 genes) compared to all genes detected as eRNA. Note that CPM values in Figure 3 differ slightly from those in Figures 2D and 2E due to different samples included in expression normalization.

GSEA showed no significant enrichment for genes specifically detected by each filter. However, GSEA revealed that genes not detected by 0.4 µm filter were enriched in pathways, such as nucleus (GO: 0005634), cellular protein modification process (GO: 0006464), nuclear protein-containing complex (GO: 0140513), transferase complex (GO: 1990234), and membrane-enclosed lumen (GO: 0031974) (Figure 3G). The enrichment of pathways such as the nucleus and membrane-enclosed lumen suggests that genes expressed within the nucleus and intracellular organelles may be released into the water, protected within membranes, and thus retained by 3 µm or 10 µm filters. These genes specific to 3 µm and 10 µm filters had lower CPM and TIN values compared to the genes detected in 0.4 µm (Figures 3B and S5).

### 3.4. Limitations and recommendations

Our findings revealed that particles in the fraction between 0.4 and 3 μm contained a higher amount of microbial RNA but a lower number of eRNA genes from Japanese medaka. Most of genes captured by 0.4 μm filter were able to be detected also by 3 μm and 10 μm filters. Therefore, we recommend the use of 3 μm filter for the non-invasive assessment of physiological status of aquatic macro-organisms. Previous laboratory experiment also concluded that 5 µm pore size cellulose membrane filter exhibited the highest efficiency of capturing microalgae eDNA and eRNA particles, in terms of time effort and filtration volume (Zaiko et al. 2022). Together with the previous finding, the use of an “appropriately” large pore size filter can increase the relative abundance of eRNA from macro-organisms and improve filtration efficiency by reducing filter clogging. This recommendation is applicable not only to RNA-Seq but also to qPCR analysis, as nuclear eRNA, in particular mRNA, may be present at much lower concentrations than eDNA (Marshall et al. 2021), making it crucial to increase the detectability. Additionally, our results showed that the genes detected in medaka eRNA overlapped with those found in skin RNA, indicating that skin is a significant source of eRNA as partly suggested by a previous study (Tsuri et al. 2021). Since skin serves as the first barrier to pathogens, eRNA may have potential as a non-invasive tool for immune assessments. Further transcriptomic comparison with other possible sources (e.g., urine, feces, gill) and from species in different taxon (e.g., Arthropod, Mollusca, Echinoderm) would be helpful in characterizing the source of eRNA.

Even using 3 μm filter and deep sequencing, the sequencing depth for the medaka transcriptome was not sufficient for quantitative analysis, as described above. Therefore, we recommend that our findings should be further developed in combination with additional approaches, such as removal of microbial RNA, PCR amplification (i.e., targeted RNA-Seq (Wang et al. 2018)), and direct RNA sequencing (e.g., adaptive sampling by Oxford Nanopore Technologies (Wang et al. 2024)), all of which would increase selectivity of RNA sequencing for target species and improve both quantitative and functional analysis. Since each of these additional approaches requires eRNA collection, our findings would remain valuable. Although requiring substantially further research, methodological advance of eRNA analysis will be essential to expand its applicability for ecological monitoring and health assessments in fields.

## ASSOCIATED CONTENT

### Supporting Information

Amount of filtered water and collected eRNA (Table S1); summary of RNA-sequencing (Table S2); results of pathway enrichment analysis (Tables S3−S6); abundance of bacteria and fungi (Figure S1); relationship between sequencing depth and the number of detected genes (Figure S2); enriched pathway networks (Figure S3); relationship between TIN and CPM in eRNA (Figure S4); distribution of TIN in eRNA (Figure S5) (PDF)

List of detected genes in skin and water (Table S7) (XLSX)

## AUTHOR INFORMATION

### Author contributions

Kyoshiro Hiki: Conceptualization, Methodology, Investigation, Formal analysis, Writing – original draft, review & editing. Toshiaki S Jo: Conceptualization, Methodology, Formal analysis, Writing – original draft, review & editing.

### Notes

The authors declare no conflicts of interest. The opinions expressed in this manuscript do not necessarily represent the official views of the authors’ affiliation.

## ACKNOWLEDGMENTS

The authors are grateful to Yukiko Ohata (National Institute for Environmental Studies) for her help in conducting molecular experiments, and to Hiroko Takahashi (National Institute for Environmental Studies) for her help in culturing test organisms. This study was financially supported by JST, CREST (Grant Number: JPMJCR22D2) and by Grant-in-Aid for JSPS Research Fellows (Grant Numbers JP22J00439 and JP22KJ3043).

## Notes

https://www.ncbi.nlm.nih.gov/bioproject/PRJDB15208

### Competing Interest Statement

The authors have declared no competing interest.

### Summary of Updates

Author name was corrected; "Toshiaki Sohma Jo" to "Toshiaki Souma Jo".

