## Supplementary Files for "Comprehensive Sequencing of Environmental RNA from Japanese Medaka at Various Size Fractions and Comparison with Skin RNA"

|  | | **Content** | **Page** |
| --- | --- | --- | --- |
| **Tables** | Table S1 | Amount of filtered water and collected eRNA | S2 |
|  | Table S2 | Summary of RNA-sequencing | S3 |
|  | Table S3 | Result of pathway enrichment analysis | S4 |
|  | Table S4 | Result of pathway enrichment analysis | S5 |
|  | Table S5 | Result of pathway enrichment analysis | S6 |
|  | Table S6 | Result of pathway enrichment analysis | S7 |
| **Figures** | Figure S1 | Relative abundance of bacteria and fungi | S8 |
|  | Figure S2 | Relationship between sequencing depth and the number of detected genes | S9 |
|  | Figure S3 | Enriched pathway networks | S10 |
|  | Figure S4 | Relationship between TIN and logarithmic CPM in eRNA | S11 |
|  | Figure S5 | Distribution of TIN in eRNA | S11 |
| **References** | | | S12 |

**Table of contents**

Table S1. Amount of filtered water and collected eRNA.

| Tank No. | Amount of filtered water (mL) | Yield of collected eRNA (ng) ^a^ | | |
| --- | --- | --- | --- | --- |
|  |  | 0.4 μm filter | 3 μm filter | 10 μm filter |
| 1 | 1250 | 955 | 154 | 279 |
| 2 | 750 | 675 | 259 | 212 |
| 3 | 750 | 780 | 234 | 237 |
| 4 | 750 | 875 | 101 | 108 |

a: RNA concentrations was estimated using 4200 TapeStation.

Table S2. Summary of RNA-sequencing.

|  | Replicate | Total reads | GC content (%) | Q30 (%) | Ratio of mapped reads (%) |
| --- | --- | --- | --- | --- | --- |
| eRNA 0.4 μm filter | 1 | 150,772,764 | 52.5 | 94.0 | 0.29 |
|  | 2 | 161,486,928 | 52.9 | 94.2 | 0.31 |
|  | 3 | 161,037,430 | 52.9 | 94.0 | 0.26 |
|  | 4 | 183,561,654 | 52.8 | 93.8 | 0.34 |
| eRNA 3 μm filter | 1 | 153,516,760 | 50.2 | 93.6 | 3.26 |
|  | 2 | 184,463,196 | 47.8 | 93.4 | 0.89 |
|  | 3 | 170,859,914 | 49.5 | 93.5 | 0.97 |
|  | 4 | 182,358,464 | 49.2 | 92.9 | 1.11 |
| eRNA 10 μm filter | 1 | 185,211,386 | 50.8 | 94.3 | 0.57 |
|  | 2 | 183,397,400 | 50.6 | 94.7 | 0.65 |
|  | 3 | 167,918,750 | 49.8 | 93.5 | 0.53 |
|  | 4 | 160,911,554 | 50.5 | 94.5 | 0.45 |
| Skin RNA | 1 | 135,617,132 | 50.6 | 93.7 | 84.1 |
|  | 2 | 156,889,058 | 50.5 | 92.7 | 76.4 |
|  | 3 | 158,187,058 | 50.7 | 93.0 | 87.5 |
|  | 4 | 158,754,782 | 50.3 | 93.1 | 75.6 |

Table S3. Top ten pathways enriched in fish skin RNA compared to the whole medaka genome.

| Enrichment false-discovery rate (FDR) corrected p-values | No. of genes | No. of total genes in pathways | Fold enrichment | Pathway |
| --- | --- | --- | --- | --- |
| < 0.001 | 1541 | 3004 | 1.5 | GO:0044267 cellular protein metabolic process |
| < 0.001 | 2043 | 4279 | 1.4 | GO:1901564 organonitrogen compound metabolic process |
| < 0.001 | 1729 | 3600 | 1.4 | GO:0019538 protein metabolic process |
| < 0.001 | 1767 | 3829 | 1.4 | GO:0044260 cellular macromolecule metabolic process |
| < 0.001 | 1547 | 3288 | 1.4 | GO:0032991 protein-containing complex |
| < 0.001 | 903 | 1709 | 1.6 | GO:0012505 endomembrane system |
| < 0.001 | 653 | 1151 | 1.7 | GO:0009056 catabolic process |
| < 0.001 | 1728 | 3802 | 1.4 | GO:0005634 nucleus |
| < 0.001 | 489 | 800 | 1.8 | GO:0003723 RNA binding |
| < 0.001 | 540 | 911 | 1.7 | GO:1902494 catalytic complex |

Table S4. Top ten pathways enriched in skin-specific genes (3987 genes) compared to all genes detected in skin RNA.

| Enrichment false-discovery rate (FDR) corrected p-values | No. of genes | No. of total genes in pathways | Fold enrichment | Pathway |
| --- | --- | --- | --- | --- |
| < 0.001 | 114 | 255 | 1.6 | GO:0034470 ncRNA processing |
| < 0.001 | 42 | 94 | 2.1 | GO:0008033 tRNA processing |
| < 0.001 | 275 | 804 | 1.3 | GO:0005739 mitochondrion |
| < 0.001 | 136 | 336 | 1.5 | GO:0034660 ncRNA metabolic process |
| < 0.001 | 167 | 1341 | 1.4 | GO:0005576 extracellular region |
| < 0.001 | 63 | 141 | 1.8 | GO:0006399 tRNA metabolic process |
| < 0.001 | 67 | 260 | 1.7 | GO:0016741 transferase activity transferring one-carbon groups |
| < 0.001 | 49 | 124 | 1.8 | GO:0009451 RNA modification |
| < 0.001 | 29 | 66 | 2.1 | GO:0006400 tRNA modification |
| < 0.001 | 113 | 937 | 1.5 | GO:0005615 extracellular space |

Table S5. Top ten pathways enriched in genes detected in both water and skin (5148 genes) compared to all genes detected in skin RNA.

| Enrichment false-discovery rate (FDR) corrected p-values | No. of genes | No. of total genes in pathways | Fold enrichment | Pathway |
| --- | --- | --- | --- | --- |
| < 0.001 | 51 | 65 | 1.7 | GO:0022626 cytosolic ribosome |
| < 0.001 | 591 | 1709 | 1.2 | GO:0012505 endomembrane system |
| < 0.001 | 151 | 288 | 1.3 | Path:ola05132 Salmonella infection |
| < 0.001 | 220 | 608 | 1.3 | GO:0005794 Golgi apparatus |
| < 0.001 | 33 | 43 | 1.8 | GO:0022625 cytosolic large ribosomal subunit |
| < 0.001 | 260 | 726 | 1.2 | GO:0031982 vesicle |
| < 0.001 | 255 | 705 | 1.2 | GO:0031410 cytoplasmic vesicle |
| < 0.001 | 255 | 705 | 1.2 | GO:0097708 intracellular vesicle |
| < 0.001 | 150 | 317 | 1.3 | Path:ola04144 Endocytosis |
| < 0.001 | 64 | 116 | 1.5 | Path:ola04070 Phosphatidylinositol signaling system |

Table S6. T Pathway networks enriched in genes that were not detected in 0.4 µm filter (5368 genes) compared to all genes detected as eRNA.

| Enrichment false-discovery rate (FDR) corrected p-values | No. of genes | No. of total genes in pathways | Fold enrichment | Pathway |
| --- | --- | --- | --- | --- |
| < 0.001 | 256 | 765 | 1.1 | GO:0140513 nuclear protein-containing complex |
| < 0.001 | 805 | 3504 | 1.1 | GO:0090304 nucleic acid metabolic process |
| < 0.001 | 727 | 2349 | 1.1 | GO:0036211 protein modification process |
| < 0.001 | 746 | 2486 | 1.1 | GO:0043412 macromolecule modification |
| 0.002 | 727 | 2348 | 1.1 | GO:0006464 cellular protein modification process |
| 0.002 | 730 | 2841 | 1.1 | GO:0016070 RNA metabolic process |
| 0.005 | 175 | 478 | 1.1 | GO:1990234 transferase complex |
| 0.005 | 578 | 2228 | 1.1 | GO:0019219 regulation of nucleobase-containing compound metabolic process |
| 0.005 | 904 | 3934 | 1.1 | GO:0046483 heterocycle metabolic process |
| 0.005 | 890 | 3845 | 1.1 | GO:0006139 nucleobase-containing compound metabolic process |

**A) Bacteria**

|  | Acidovorax | Aeromonas | Collimonas | Escherichia | Herbaspirillum | Janthinobacterium | Massilia | Noviherbaspirillum | Pseudomonas | Shewanella | Shigella | Others |
| --- | --- | --- | --- | --- | --- | --- | --- | --- | --- | --- | --- | --- |
| 0.4 µm-1 | 1.11719 | 1.70999 | 6.40675 | 0.889193 | 23.4838 | 4.83356 | 21.9334 | 14.7515 | 0 | 0 | 0.159599 | 24.715 |
| 0.4 µm-2 | 2.6087 | 3.53623 | 6.70531 | 0.405797 | 21.314 | 4.79227 | 20.1546 | 14.2609 | 0 | 0.463768 | 0 | 25.7585 |
| 0.4 µm-3 | 4.5236 | 3.05944 | 5.28846 | 1.0271 | 18.8811 | 3.93357 | 16.4773 | 13.3741 | 1.09266 | 1.0271 | 0.262238 | 31.0533 |
| 0.4 µm-4 | 1.81306 | 1.4082 | 6.33691 | 0.404858 | 22.4256 | 5.10474 | 20.8766 | 14.0644 | 0.633691 | 0.281641 | 0.14082 | 26.5094 |
| 3 µm-1 | 1.51845 | 69.4192 | 0.284332 | 0.30853 | 1.07683 | 0.187538 | 0.798548 | 0.961887 | 0.544465 | 5.95886 | 0.0665457 | 18.8748 |
| 3 µm-2 | 2.51116 | 60.119 | 0.297619 | 3.32961 | 1.07887 | 0 | 0.558036 | 0.948661 | 0.63244 | 4.81771 | 0.558036 | 25.1488 |
| 3 µm-3 | 2.56757 | 48.7725 | 0.563063 | 0.641892 | 2.38739 | 0.495495 | 1.66667 | 1.89189 | 8.09685 | 4.56081 | 0.202703 | 28.1532 |
| 3 µm-4 | 1.86804 | 54.4515 | 0.417329 | 0.808161 | 1.62295 | 0.317965 | 1.29173 | 1.11288 | 7.92925 | 4.65686 | 0.178855 | 25.3445 |
| 10 µm-1 | 2.5641 | 60.5389 | 0.65189 | 0.956106 | 1.78183 | 0.391134 | 1.43416 | 1.1734 | 1.3907 | 6.38853 | 0 | 22.7292 |
| 10 µm-2 | 1.68934 | 60.9959 | 0.304883 | 0.737251 | 1.08608 | 0.209484 | 0.841891 | 0.805644 | 6.31405 | 6.41804 | 0.109133 | 20.4883 |
| 10 µm-3 | 0.822095 | 68.16 | 0 | 0.175503 | 0.230926 | 0 | 0.147792 | 0.18474 | 8.31332 | 7.5836 | 0 | 14.382 |
| 10 µm-4 | 1.50311 | 60.8332 | 0.150311 | 1.00923 | 0.708611 | 0.128838 | 0.49388 | 0.751557 | 7.62293 | 7.04316 | 0.257677 | 19.4975 |
| Skin-1 | 0.737429 | 8.32373 | 0 | 81.1587 | 0.0460893 | 0 | 0.0599161 | 1.80209 | 0.677513 | 0.0368715 | 0.700558 | 6.45711 |
| Skin-2 | 1.29945 | 1.11157 | 0.0574053 | 84.1457 | 0.360088 | 0 | 0.0260933 | 0.918485 | 0.0991546 | 0 | 5.91274 | 6.0693 |
| Skin-3 | 0.810127 | 4.89971 | 0 | 86.3174 | 0 | 0 | 0 | 2.1149 | 0.12853 | 0 | 0.163583 | 5.56573 |
| Skin-4 | 1.42594 | 3.36362 | 0.0632475 | 80.2208 | 0.471481 | 0 | 0.0402484 | 0.799218 | 0.592226 | 0.0402484 | 7.12971 | 5.85327 |

**B) Fungi**

|  | Aspergillus | Biomphalaria | Candida | Colletotrichum | Cyphellophora | Enterocytozoon | Lobosporangium | Malassezia | Mitosporidium | Rhodotorula | Saccharomyces | Sordaria | Torulaspora | Wickerhamomyces |
| --- | --- | --- | --- | --- | --- | --- | --- | --- | --- | --- | --- | --- | --- | --- |
| 0.4 µm-1 | 9.69388 | 0 | 0 | 7.14286 | 0 | 0 | 5.10204 | 0 | 6.63265 | 0 | 0 | 21.4286 | 0 | 50 |
| 0.4 µm-2 | 4.36364 | 2.90909 | 0 | 5.81818 | 0 | 3.27273 | 1.81818 | 2.90909 | 15.2727 | 0 | 2.90909 | 14.5455 | 0 | 46.1818 |
| 0.4 µm-3 | 12.0805 | 0 | 0 | 4.36242 | 0 | 9.73154 | 0 | 0 | 6.04027 | 0 | 10.0671 | 13.0872 | 0 | 44.6309 |
| 0.4 µm-4 | 21.9124 | 0 | 9.16335 | 0 | 0 | 4.38247 | 0 | 1.99203 | 0 | 0 | 24.7012 | 24.3028 | 0 | 13.5458 |
| 3 µm-1 | 2.71186 | 0 | 0 | 3.05085 | 4.0678 | 0 | 4.23729 | 7.11864 | 7.62712 | 0 | 32.8814 | 8.47458 | 0 | 29.8305 |
| 3 µm-2 | 13.41 | 0 | 0 | 12.6437 | 0 | 2.29885 | 8.04598 | 5.74713 | 8.42912 | 0 | 12.6437 | 19.1571 | 2.29885 | 15.3257 |
| 3 µm-3 | 7.57946 | 0 | 0 | 14.9144 | 0 | 5.62347 | 1.46699 | 5.86797 | 4.88998 | 5.62347 | 13.6919 | 14.1809 | 0 | 26.1614 |
| 3 µm-4 | 6.07477 | 0 | 0 | 18.2243 | 3.03738 | 5.14019 | 5.14019 | 2.1028 | 6.54206 | 0 | 10.9813 | 15.6542 | 0 | 27.1028 |
| 10 µm-1 | 8.21256 | 0 | 0 | 12.5604 | 0 | 0 | 0 | 0 | 3.86473 | 0 | 26.087 | 37.6812 | 0 | 11.5942 |
| 10 µm-2 | 21.2389 | 6.19469 | 0 | 5.30973 | 0 | 0 | 7.9646 | 0 | 9.73451 | 0 | 6.19469 | 29.2035 | 0 | 14.1593 |
| 10 µm-3 | 15.6863 | 0 | 0 | 18.3007 | 0 | 3.92157 | 5.88235 | 0 | 10.4575 | 0 | 0 | 30.719 | 0 | 15.0327 |
| 10 µm-4 | 11.6438 | 0 | 0 | 13.6986 | 0 | 3.42466 | 7.53425 | 0 | 4.79452 | 0 | 6.84932 | 30.137 | 0 | 21.9178 |
| Skin-1 | 21.9388 | 0 | 5.61224 | 0 | 0 | 5.10204 | 0 | 7.65306 | 0 | 0 | 16.3265 | 25 | 0 | 18.3673 |
| Skin-2 | 31.1005 | 0 | 6.69856 | 0 | 0 | 6.69856 | 0 | 0 | 0 | 0 | 0 | 30.622 | 0 | 24.8804 |
| Skin-3 | 16.6667 | 0 | 3.64583 | 0 | 0 | 6.77083 | 0 | 6.77083 | 0 | 0 | 21.875 | 27.0833 | 0 | 17.1875 |
| Skin-4 | 23.2353 | 0 | 5.31888 | 0 | 0 | 6.19048 | 0 | 4.80796 | 0 | 0 | 12.7338 | 27.5684 | 0 | 20.1451 |

Figure S1. Relative abundance of (A) bacteria and (B) fungi at genus levels in water and fish skin RNA samples. Taxonomic identification was performed using DecontaMiner (Sangiovanni et al. 2019). While the fungal genus composition showed little variation among sample types, bacterial composition differed significantly. For example, *Escherichia* dominated skin RNA (> 80%) but was less than 4% in water. The genus *Herbaspirillum, Massilia*, and *Noviherbaspirillum* were abundant (> 10%) in water collected by the 0.4 µm filter, whereas their proportions were below 2.5% in water collected by the 3 and 10 µm filters.

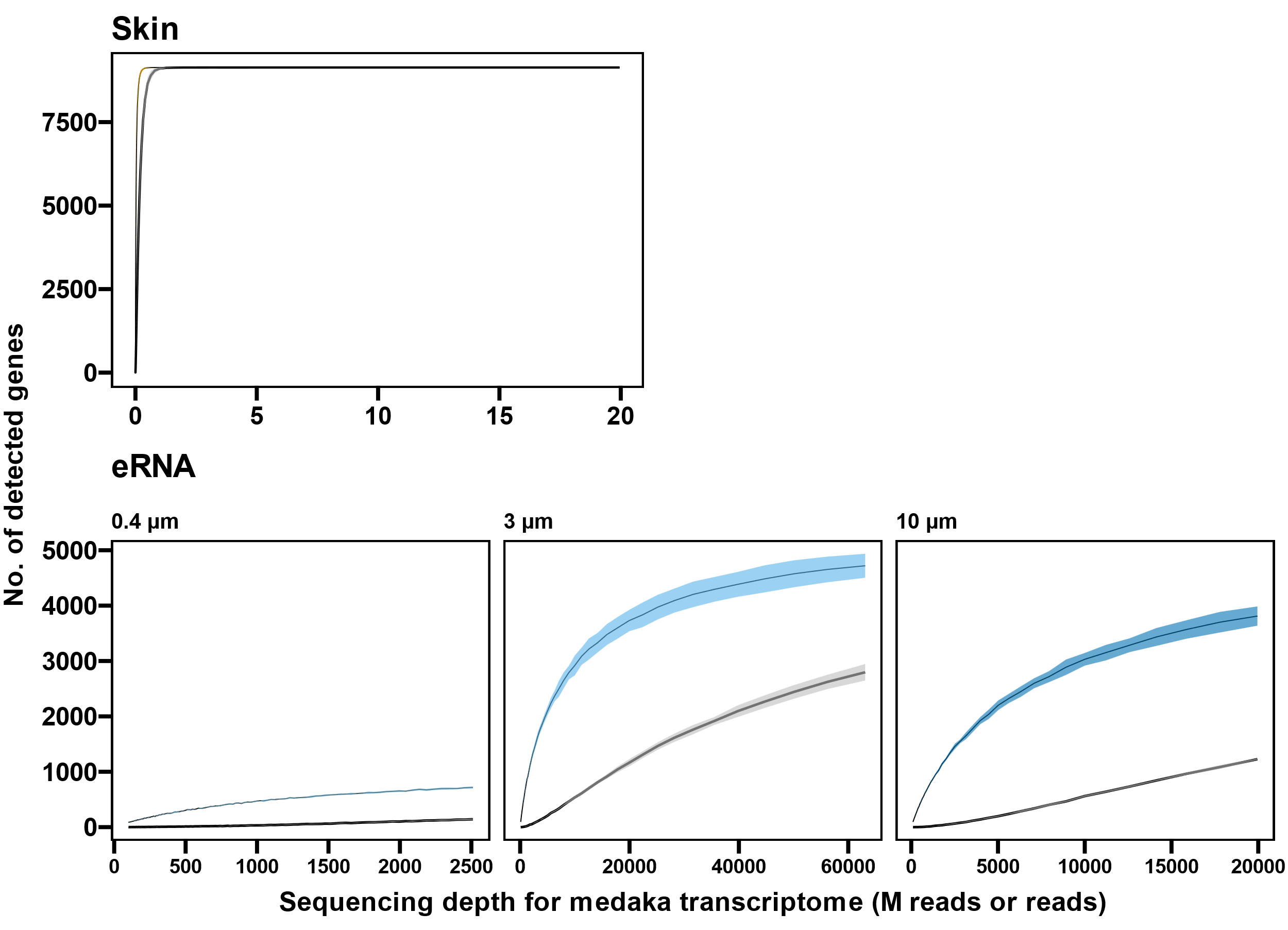

Figure S2. The number of detected genes as a function of the sequencing depth for medaka transcriptome (million reads for skin RNA and reads for eRNA). Curves and shaded areas represent the mean and standard deviations (*n* = 4), respectively, both of which were calculated by Monte Carlo simulations. Colored and gray lines indicate all genes and genes with > 5 read counts, respectively. Note that the x-axis indicates the sequencing depth mapped to the medaka transcriptome rather than the entirety of bulk RNA, which is more than 30−100 times greater for eRNA than the values shown.

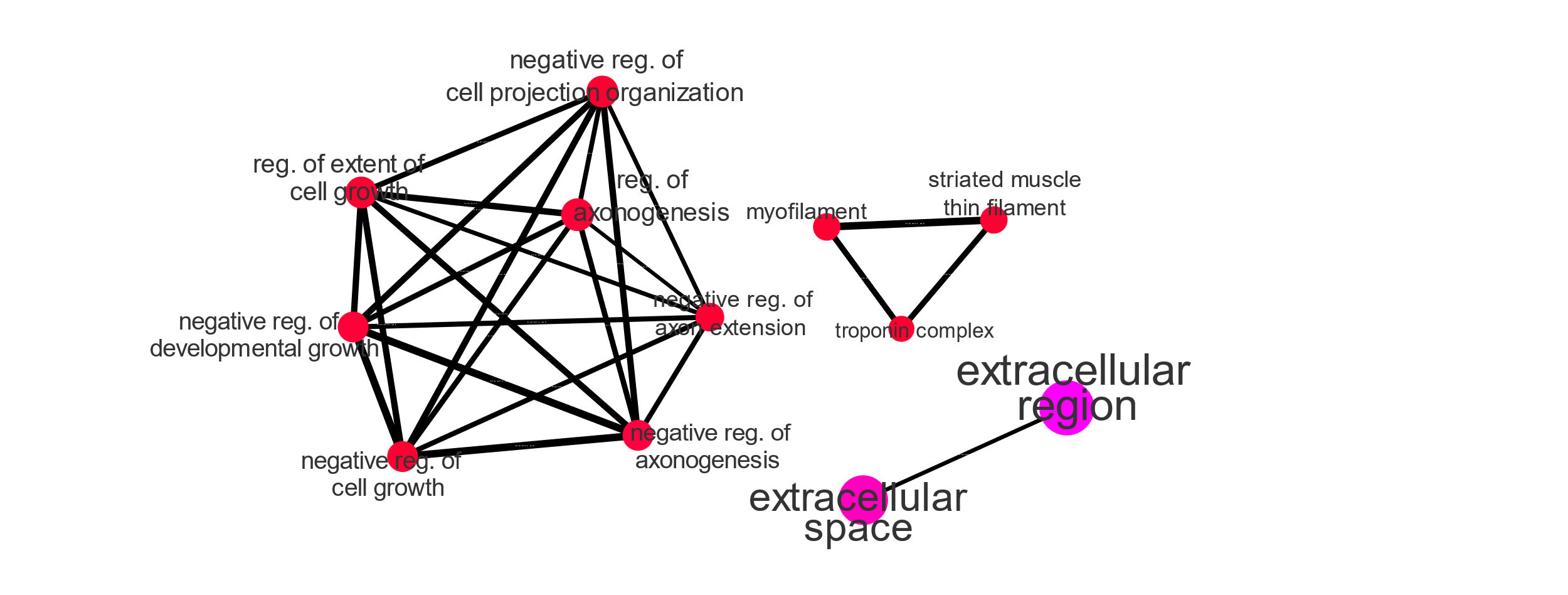

Figure S3. Pathway networks enriched in eRNA-specific RNA (1192 genes) compared to all genes detected in eRNA. Vivid color and larger nodes are more significantly enriched (FDR-adjusted p-value < 0.05) and larger gene sets, respectively. Edge widths represent the degree of gene overlap. Enrichment analysis was performed using ShinyGO ver. 0.80 (Ge et al. 2020). The pathways related to negative regulation of axonogenesis included semaphorin family genes, while the pathways related to myofilament included troponin family genes.

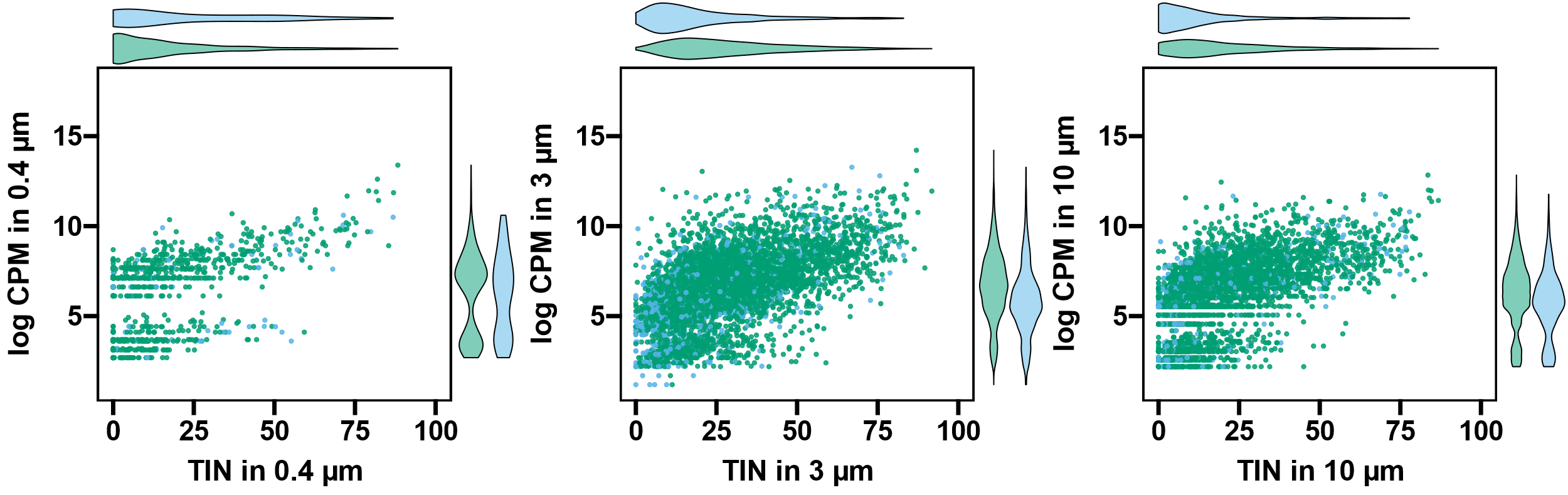

Figure S4. Relation between transcript integrity number (TIN) and logarithmic CPM in eRNA collected using (Left) the 0.4 µm, (Center) 3 µm, and (Right) 10 µm filters. The center panel is identical to Figure 2D. Green indicates genes detected in both eRNA and fish skin RNA, while light blue represents genes specifically detected in eRNA, respectively.

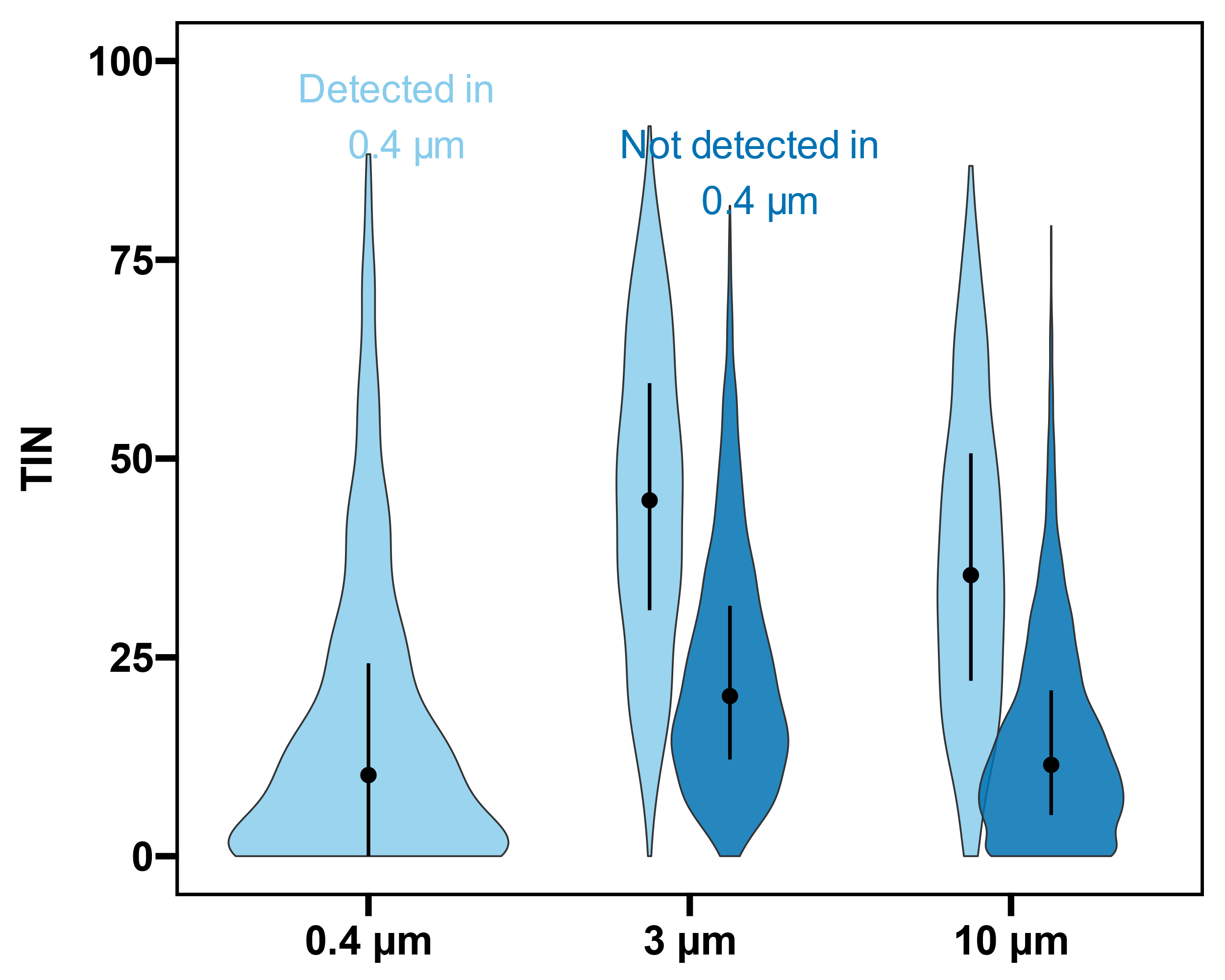

Figure S5. Violin plots showing the distribution of TIN for eRNA across different filters. Error bars indicate the median and interquartile range (25 and 75 percentiles). Light blue represents genes detected in the 0.4 µm filter, while blue represents genes not detected in the 0.4 µm filter.
